# SCAN-ATAC-Sim: a scalable and efficient method for simulating single-cell ATAC-seq data from bulk-tissue experiments

**DOI:** 10.1101/2020.05.29.123638

**Authors:** Zhanlin Chen, Jing Zhang, Jason Liu, Zixuan Zhang, Jiangqi Zhu, Donghoon Lee, Min Xu, Mark Gerstein

## Abstract

**Summary:** scATAC-seq is a powerful approach for characterizing cell-type-specific regulatory landscapes. However, it is difficult to benchmark the performance of various scATAC-seq analysis techniques (such as clustering and deconvolution) without having *a priori* a known set of gold-standard cell types. To simulate scATAC-seq experiments with known cell-type labels, we introduce an efficient and scalable scATAC-seq simulation method (SCAN-ATAC-Sim) that down-samples bulk ATAC-seq data (e.g., from representative cell lines or tissues). Our protocol uses a consistent but tunable signal-to-noise ratio across cell types in a scATAC-seq simulation for integrating bulk experiments with different levels of background noise, and it independently samples twice without replacement to account for the diploid genome. Because it uses an efficient weighted reservoir sampling algorithm and is highly parallelizable with OpenMP, our implementation in C++ allows millions of cells to be simulated in less than an hour on a laptop computer.

*Availability:* SCAN-ATAC-Sim is available at scan-atac-sim.gersteinlab.org.

*Contact:*

**Supplementary information:** Supplementary data are available at *Bioinformatics* online.

## Introduction

High-resolution single-cell assay for transposase-accessible chromatin using sequencing (scATAC-seq) techniques reveal transcriptional landscapes in a cell-type-specific manner (Buenrostro, et al., 2015). Numerous scATAC-seq data analysis approaches, such as those employed for calling and defining active regions in various cell types, clustering, and deconvolution, have been published (Bravo Gonzalez-Blas, et al., 2019; Fang, et al., 2019; Liu, et al., 2019; Schep, et al., 2017; Xiong, et al., 2019; Zamanighomi, et al., 2018). However, it has been difficult to evaluate the efficacy of these techniques because we do not have *a priori* knowledge of gold-standard cell types. One way to evaluate these analysis methods is to simulate scATAC-seq with ground-truth labels. With this approach, the analysis methods can be benchmarked against one another with quantifiable parameters that affect the separability of different cell types.

There are three major challenges in simulating realistic scATAC-seq data. First, each open chromatin region can only be captured zero, one, or two times in a diploid genome, resulting in at most two reads at one locus in scATAC-seq. Second, similar to bulk ATAC-seq data, many reads in scATAC-seq come from non-peak, background regions. Third, it is computationally expensive to simulate a dataset with millions of cells in order to evaluate the performance of an analysis method on large datasets. Currently, there are two existing approaches for simulating scATAC-seq data, and both approaches are limited (Table 1).

**Table 1.** Comparison between SCAN-ATAC Sim and previous methods and a brief list of parameters that control the simulation.

| Features | Direct Down Sampling | Peak Region Sampling | SCAN-ATAC Sim |
| --- | --- | --- | --- |
| Read-level simulation | Yes | No | Yes |
| Flexible signal-to-noise | No | No | Yes |
| Diploid genomic constraint | No | Yes | Yes |
| Short runtime | No | No | Yes |

| Long Flag | Default Value | Long Flag | Default Value |
| --- | --- | --- | --- |
| --cell_number | 10,000 | --min_frag | 1,000 |
| --signal_to_noise | 0.7 | --max_frag | 20,000 |
| --frag_num | 3,000 | --extend_peak_size | 1,000 |
| --variance | 0.5 | --bin_size | 1,000 |

The first approach randomly samples reads from a curated set of bulk ATAC-seq data (Fang, et al., 2019). Bulk ATAC-seq has dramatic variations in the signal-to-noise ratio between experiments. However, cells in scATAC-seq experiments undergo similar procedures, resulting inless background variation. Directly sampling the bulk ATAC-seq can introduce cell-type-specific backgrounds such as bias from batch effects, confounding downstream analyses. Moreover, it may extract more than two reads for a single genomic locus, thus violating the diploid nature of scATAC-seq experiments. Lastly, direct down-sampling is inefficient. We replicated the method, and it used 26.6 hours to simulate one million cells.

The second method simulates individual cells by selecting foreground peaks with a strictly fixed signal-to-noise ratio (Xiong, et al., 2019; Zhang, et al., 2020). In reality, over half of the reads in scATAC-seq come from background regions. Even though it utilizes drop-out ratios and adheres to the diploid constraint, peak region sampling ignores the background; thus, it is limited in representing real data. Further, this method does not simulate reads. Rather, each cell is represented by simulated foreground peaks. Analysis methods that construct a cell-by-bin read coverage matrix cannot be evaluated using a dataset simulated from this method due to the lack of read-level information. Lastly, repeated sampling from a binomial distribution for all peak regions also limits the efficiency when scaled to millions of cells.

We aim to address these challenges. The lack of a standard, representational synthetic scATAC-seq dataset motivated SCAN-ATAC-Sim, which offers an improvement in simulation quality and reduction in runtime compared to both previous approaches. Our command line software, implemented in C++ with OpenMP parallelization, takes BAM files from bulk ATAC-seq experiments as input and outputs sampled reads for each cell based on user-provided parameters such as the number of reads per cell, total cell number, and signal-to-noise ratio (Table 1).

## 2 Methods

SCAN-ATAC Sim consists of two main steps: data preprocessing and single-cell simulation. Briefly, the process starts with BAM files of bulk ATAC-seq experiments for desired cell types (Fig. 1a). The data preprocessing step defines a cell-type-specific foreground from the merged peaks and a unified background (Fig. 1b). For each cell, the single-cell simulation step samples the foreground and background regions twice without replacement with the probability proportional to the read coverage (Fig. 1c) and randomly selects one read for each sampled region (Fig. 1d). Then, reads from both the foreground and background are combined to form reads in one cell (Fig. 1e). This single-cell simulation step is then repeated for a large number of cells as specified by the user-provided parameter.

**Figure 1.**
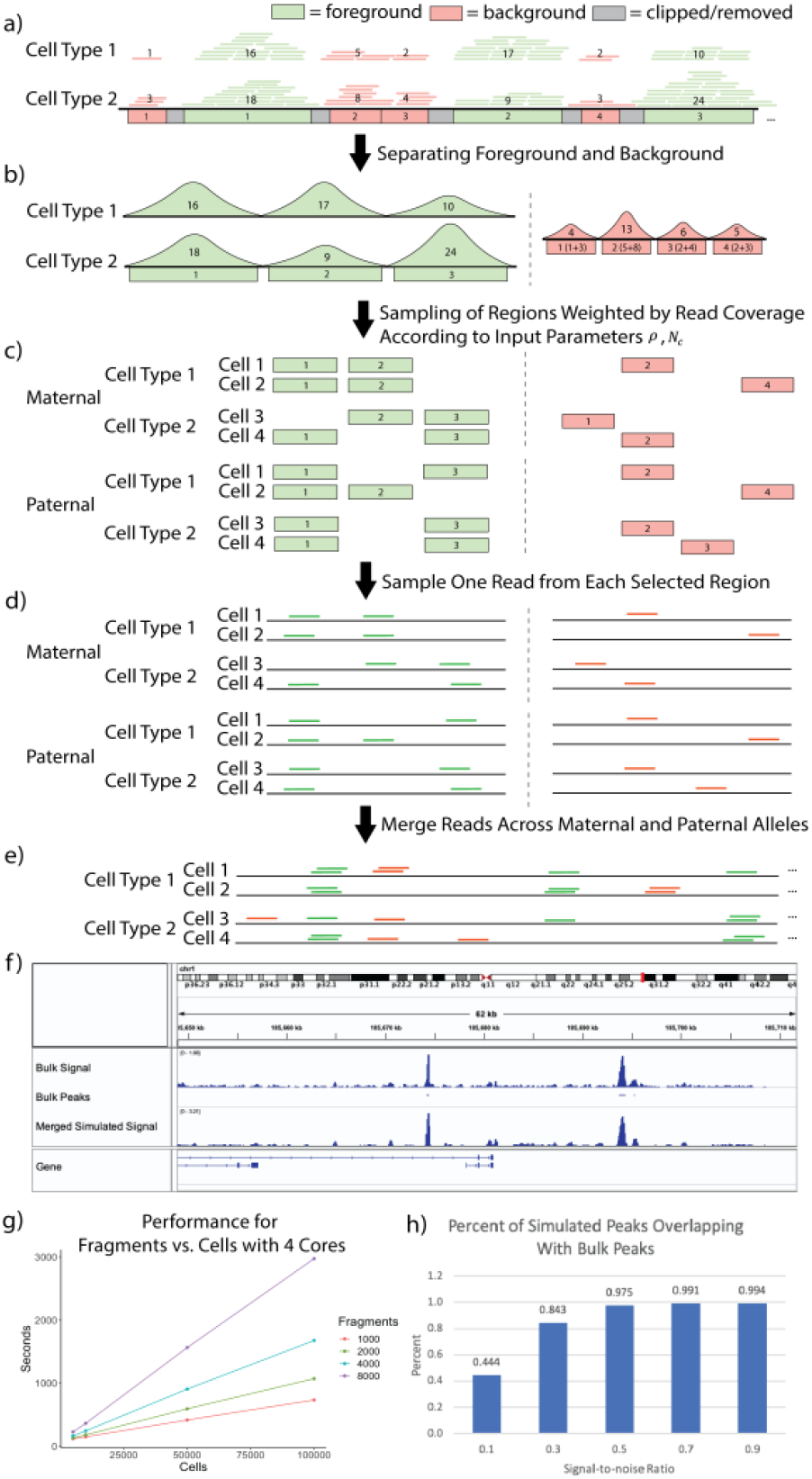
a) Bulk-tissue ATAC-seq reads are partitioned into foreground and background based on overlap with merged peaks. The number in each region indicates the read coverage. b) The cell-type-specific foreground reads are separated, and a unified background is created by combining background reads across all cell lines. c) The regions are sampled without replacement, with the read coverage as weights for the foreground and background, for paternal and maternal alleles. d) Reads are sampled from the selected regions using a uniform distribution. e) All sampled reads are combined to form reads covering a cell. f) chr1 visualization of bulk and simulated (*ρ* = 0.4, *f* = 1*k,c* = 100*k*) CLP cells is shown in the Integrative Genome Browser. g) Performance for region number *N_c_* vs. cell number is shown for four cores. h) Percentage of peaks from simulated CLP cells that overlap with CLP bulk peaks is shown, demonstrating the relationship between the signal-to-noise ratio and the cell-type specificity of the simulation.

### 2.1 Data Preprocessing

The foreground regions are defined by merging peaks from various cell types. Then, the background regions are defined as the complement of the 1 kb-extended foreground divided into bins of fixed sizes (Fig. 1a). The paired and de-duplicated reads from each cell type are intersected with the foreground region to obtain cell-type-specific foreground reads, and unified background weights are created by combining background reads from all cell types (Fig. 1b).

### 2.2 Single-Cell Simulation

First, for a cell of cell type *c*, the parameter *N_c_* determines the total number of reads (or regions, since we only sample one read from each region) in an individual cell. *N_c_* can vary with a log-normal distribution so that every cell has a slightly different number of reads. We also designate a user parameter for the signal-to-noise ratio *ρ*. We use *N_c_* and *ρ* to calculate the allocation of foreground and background reads in a cell so that 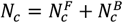 (1).

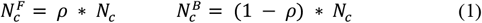

A high signal-to-noise ratio will allocate more reads for foreground regions, thereby making the cell types more separable. Once an allocation between foreground and background is made, the allocations are further halved to mimic reads coming from maternal 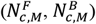 and paternal 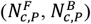 alleles. Next, SCAN-ATAC Sim performs an efficient two-step sampling to generate representative regions. First, for each cell, 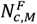 and 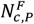 regions are separately sampled in two independent trials. Each trial samples without replacement and uses read coverage from the corresponding cell type as weights (Fig. 1c). Representative background regions are generated in a similar manner, with averaged weights from all cell types. Second, one read is randomly sampled from each selected region with equal probability (Fig. 1d). Hence, for any given region in the genome, we can guarantee that no more than two reads are sampled from the maternal and paternal trials because each trial samples without replacement. Lastly, the foreground and background reads are merged for a total of 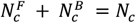 reads in one cell (Fig. 1e). This process is repeated many times to simulate a massive number of cells.

Specifically, if 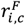 represents the total number of reads in the *i^th^* foreground region for cell type *c*, then the probability of sampling region *i* can be calculated as 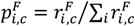. In contrast, we calculate a uniform background sampling probability for the background region. For instance, if 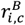 represents the read count in the *i^th^* background region for cell type *c*, then the uniform background sampling rate for the *i^th^* background region is 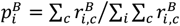.

## 3 Results

### 3.1 Comparing Simulated scATAC-seq with Bulk ATAC-seq

We analyzed how the simulation mimics cell-type specificity. We simulated and aggregated common lymphoid progenitor (CLP) cells under various signal-to-noise ratios. The pileup signal distribution of simulated scATAC-seq was similar to bulk ATAC-seq (Fig. 1f). We also called peaks from the bulk and simulated data. The amount of overlap between simulated and bulk peaks increased as the signal-to-noise ratio increased (Fig. 1h), indicating that the signal-to-noise ratio contributed to the celltype specificity of chromatin accessibility, which can be used to vary the difficulty of the benchmarking task. Furthermore, the simulation retained cell-type specificity across different cell types (Fig. S2).

### 3.2 Analyzing Simulated scATAC-seq Data

Next, we analyzed the simulated scATAC-seq data using SnapATAC (Fang, et al., 2019). Under high signal-to-noise ratios, SnapATAC clustered and labeled cells with high accuracy (Fig. S3). As the signal-to-noise ratio decreased, the separability between certain cell types decreased in the cell clustering. The read coverage also influenced the analysis outcome, with more reads per cell increasing the foreground signal and separability (Fig. S4).

### 3.3 Complexity and Runtime

There are two major computational challenges to simulating scATAC-seq data. First, up to millions of representative foreground and background regions can be selected with varying probabilities without replacement for one cell. Second, one experiment can contain tens of thousands to millions of cells. To address these challenges, we used two techniques to implement a highly efficient and scalable software for the sampling procedures mentioned in Section 2.3.

First, to improve the efficiency of single-cell sampling, we implemented a reservoir-sampling algorithm to select the representative regions. If *n* represents the number of regions and *m* represents the number of regions to be selected, weighted reservoir sampling without replacement can be performed with *O*(*n*) + *O*(*m · log*(*n/m*))*O*(*logm*), as compared to *O*(*n·m*) for traditional weighted sampling methods (Efraimidis and Spirakis, 2006). Especially in our case where *n ≫ m*, weighted reservoir sampling approaches *O*(*n*).

Second, we further parallelized our method in a cell-wise fashion with multicore parallelism using OpenMP, which offers a scalable improvement in runtime. By utilizing both approaches, we enabled the sampling of millions of cells in less than an hour. We demonstrate that SCAN-ATAC-Sim achieves a scalable speed-up for a cell group of four cell types. The runtime does not change with signal-to-noise ratio *ρ* but varies linearly with both cell and region number *N_c_* (Fig. 1g).

## 4 Discussion

We introduce a new software tool for constructing a labeled scATAC-seq dataset from bulk ATAC-seq data via guided down-sampling. This tool is useful for evaluating the efficacy of single-cell data analysis techniques by simulating scATAC-seq data while adhering to various biological constraints. The acceleration obtained via multicore parallelism permits the simulation of millions of cells in less than an hour. Moreover, this tool is highly scalable and offers a space-time tradeoff to match the rate of growth in the number of cells sequenced for scATAC-seq.

