## Supplement for "SCAN-ATAC-Sim: a scalable and efficient method for simulating single-cell ATAC-seq data from bulk-tissue experiments" (from another version of this preprint: version 1, 2020-05-30)

### 1 Cell Group Selection

Bulk-tissue ATAC-seq for Natural Killer (NK), Common Lymphoid Precursor (CLP), Erythrocytes (ERY-1), megakaryocyte–erythroid progenitor (MEP), and Monocytes (Mono) were used as a cell group. The selected cell group is meant to resemble the different cell types in real scATAC-seq experiments. The scATAC-seq experiments simulated by our software would inherit these bulk-tissue cell line labels. For benchmarking, only scATAC-seq experiment of the NK cells were generated, so the other cell lines only contribute to the background of the simulation.

### 2 Input and Output

The input to our software is a collection of aligned bam files of the cell group. The data preprocessing step provides an intermediate output. The intermediate output consists of the read coverage of the foreground and background region and the reads intersecting with the foreground and background region for one cell line. The single-cell simulation step then uses the intermediate output to perform the simulation for each cell. The final output is foreground and background reads of every cell in the scATAC-seq experiment for one cell type.

### 3 Data Preprocessing

#### 3.1 *Foreground*

Peaks for each cell were called using MACS2 from aligned bam files using the ENCODE ATAC-seq pipeline (Zhang, et al., 2008). The narrow peaks were merged to define the foreground region.

#### 3.2 *Background*

The complement of the 1kb-extended foreground divided in bins of fixed sizes (i.e. 500bp or 1kb) were defined as the background. Hence, to make the background regions, windows of fixed sizes (i.e. 500bp or 1kb) were generated based on the hg38 genome from bedtools makewindows, and the windows that does not overlap with the 1kb-extended foreground regions were considered background regions.

#### 3.3 *Read Filtering and Processing*

Paired and deduplicated reads for each bulk-tissue ATAC-seq were filtered using the SAMtools (Li, et al., 2009) with Pysam. The filtered reads for each cell-type were then intersected with the foreground regions to obtain the foreground reads using bedtools intersect. The background reads are obtained by intersecting the midpoint of all reads with background regions to prevent overlap between multiple background regions. The intersection was also performed with bedtools intersect using pybedtools. Lastly, this step would output the background read coverage and region-to-read intersection summed across all cell types, but the foreground read coverage and region-to-read intersection is unique for each cell line. The read coverage was obtained using bedtools coverage, and the region-to-read intersection was obtained using bedtools intersect.

### 4 Single-Cell Simulation

#### 4.1 *Sampling Regions According to Read Coverage*

A total of $N_{c}$ regions are sampled for each cell. The number of regions/reads for each cell could vary across cells in single-cell sequencing, and the user can provide the mean and variance for a log-normal distribution for the region number in each cell.

The regions are sampled with read coverage as weights. The foreground and background regions are sampled separately, hence the total allocation of $N_{c}$ regions is divided into $N_{c}^{F}$ and $N_{c}^{B}$ by the signal-to-noise ratio $\rho$. Furthermore, each allele in the diploid genome could only be sampled twice or less. Therefore, the number of regions for each cell is evenly distributed across the two (maternal and paternal) rounds for the foreground and background ($N_{c,M}^{F}$ and $N_{c,P}^{F}$; $N_{c,M}^{B}$ and $N_{c,P}^{B}$).

The following series of steps are performed to sample regions represented by reads in each cell. First, the application samples $N_{c,M}^{F}$ foreground regions without replacement. After that, the application samples $N_{c,P}^{F}$ regions again from a fresh set of foregrounds without replacement. Then, it samples $N_{c,M}^{B}$ background regions without replacement. Finally, the application samples $N_{c,B}^{F}$ regions again from a fresh set of backgrounds without replacement.

Here, we make the assumption that the sampling of two alleles (maternal and paternal) on the same locus is independent of one another because a flow cell could sequence hundreds of millions of reads at the same time after amplification, effectively sampling the alleles instantaneously and independently. Previous research on single-cell allelic coverage also supports our assumption (Zhang, et al., 2015).

Weighted reservoir sampling is a novel, efficient algorithm for weighted sampling without replacement. The normalized read coverage of the regions is used as weights for weighted reservoir sampling without replacement. The main computational cost is iterating through all the regions when simulating every cell. However, exponential jumps accelerate the computation by reducing the reservoir update to only $O(n\cdot log(m))$ where n is the total population of regions and m is the region number. The algorithm also uses a heap to efficiently updates to the reservoir. Since sampling without replacement is a sequential and memory-intensive process, parallelism within each cell is very limited. However, each cell is independent of one another, so light-weight shared memory parallelism across multiple central processing units (CPU) cores with OpenMP achieves a scalable speed up (Dagum and Menon, 1998).

#### 4.2 *Sampling Reads from Each Selected Region*

Once a set of regions is obtained for each cell, a read is sampled from all reads that intersect with a region under a uniform distribution using the region-to-read intersection, which is an intermediate output from the data preprocessing step. To sample a read for each region, the application uses the region-to-read intersection to construct a dictionary that maps from one region to its intersecting reads. It counts the number of intersecting reads, then it generates a random number between zero and that read count. The number generated is the index of the selected read, and the read is accessed from the dictionary in $O(1)$ time.

The majority of the computational overhead of this step consists of creating the mapping between regions and reads, so the mapping is shared among multiple cores when parallelized with OpenMP.

### 5  Downloading and Using the SCAN-ATAC Sim software

The source code and the binary executables are available for download at [scan-atac-sim.gersteinlab.org](http://scan-atac-sim.gersteinlab.org/).

The data preprocessing step could be performed with our python script:

python SCAN-ATAC_Preprocessing.py -c <comma separated file prefixes for each cell type> -i <directory containing the peak files with prefix and .narrowPeak suffix> -j <directory containing the bam files with prefix and .bam suffix> -e <custom configuration for bp length separating between foreground and background fragments> -b <bp length for background fragment size> -o <output directory>

The dependencies for the data processing python script include python version 3.6, Pysam version 0.15.4, numpy version 1.16.4, pandas version 1.0.3, pyBigWig version 0.3.17, wget version 3.2, and pybedtools version 0.8.0. The source code could be found at <http://www.scan-atac-sim.gersteinlab.org/#/Preprocess>.

The single-cell simulation step is divided into two parts. The sampling of regions using weighted reservoir sampling is performed with the script weighted_sampling, and the sampling of reads with a uniform distribution is performed with the script uniform_sampling. Both scripts and their source code could be found at <http://www.scan-atac-sim.gersteinlab.org/#/Simulation>.

./weighted_sampling -f NKcell.peak_counts.bed -b bg_counts.bed -of NKcell.foreground.sampled.bed -ob NKcell.background.sampled.bed -n 2000 -nv 0.5 -c 10000 -s 0.6 -min 1000 -max 20000

The previous example for weighted reservoir sampling would sample the foreground and background regions of the NK cells using read coverage from data preprocessing. The number of regions for each cell follows a log-normal distribution with a mean of 2000 as specified by the -n flag and a variance of 0.5 as specified by the -nv flag, with a minimum (-min) of 1,000 and a maximum (-max) of 20,000. The number of cells simulated is 10,000 with the -c parameter, and the signal to noise is 0.6 with the -s flag. The output of the foreground and background regions are specified by the -of and -ob flag, respectively. The script automatically detects for the number of cores available for parallelization, but could also be manually set using environment variables, i.e. OMP_NUM_THREADS=4 would limit to 4 cores.

./uniform_sampling NKcell.peak_intersect.bed NKcell.foreground.sampled.bed NKcells.foreground.bed

./uniform_sampling bg_intersect.expanded.bed NKcell.background.sampled.bed NKcells.background.bed

cat NKcells.foreground.bed NKcells.background.bed > NKcells.bed

The foreground and background reads are sampled for NK cells with the selected regions and the region-to-read intersection. Lastly, the foreground and background reads are combined into a single file. The output file would contain reads of all cells from that cell line.

The process is then repeated for other cell lines in the cell group to obtain scATAC-seq experiments with different cell types.
